# Mapping and DNA sequence characterisation of the *Ry*_*sto*_ locus conferring extreme virus resistance to potato cultivar ‘White Lady’

**DOI:** 10.1101/809152

**Authors:** Mihály Kondrák, Andrea Kopp, Csilla Uri, Anita Sós-Hegedűs, Edina Csákvári, Mátyás Schiller, Endre Barta, István Cernák, Zsolt Polgár, János Taller, Zsófia Bánfalvi

## Abstract

Virus resistance genes carried by wild plant species are valuable resources for plant breeding. The *Ry*_*sto*_ gene, conferring a broad spectrum of durable resistance, originated from *Solanum stoloniferum* and was introgressed into several commercial potato cultivars, including ‘White Lady’, by classical breeding. *Ry*_*sto*_ was mapped to chromosome XII in potato, and markers used for marker-assisted selection in breeding programmes were identified. Nevertheless, there was no information on the identity of the *Ry*_*sto*_ gene. To begin to reveal the identification of *Ry*_*sto*_, fine-scale genetic mapping was performed which, in combination with chromosome walking, narrowed down the locus of the gene to approximately 1 Mb. DNA sequence analysis of the locus identified six full-length *NBS-LRR*-type (short *NLR*-type) putative resistance genes. Two of them, designated *TMV2* and *TMV3*, were similar to a *TMV resistance* gene isolated from tobacco and to *Y-1*, which co-segregates with *Ry*_*adg*_, the extreme virus resistance gene originated from *Solanum andigena* and localised to chromosome XI. Furthermore, *TMV2* of ‘White Lady’ was found to be 95% identical at the genomic sequence level with the recently isolated *Ry*_*sto*_ gene of the potato cultivar ‘Alicja’. In addition to the markers identified earlier, this work generated five tightly linked new markers which can serve potato breeding efforts for extreme virus resistance.

## Introduction

For sustainable intensification of crop production, disease control should, when possible, be achieved using genetics rather than using costly recurrent chemical sprays. Wild relatives of crop plants are a good source of genes for disease resistance. Potato (*Solanum tuberosum*), the world’s fourth most important food crop, following maize, wheat and rice, can be crossed with a number of wild *Solanum* species. Nevertheless, classical breeding for resistance is time-consuming, and it is extremely difficult to recover the parental combination necessary for beneficial alleles in the progeny. Thus, there is great value in genetic approaches that can improve disease resistance in potato varieties without disrupting favourable combinations of alleles [1].

One of the major factors adversely affecting potato production worldwide is virus infection. Viruses such as *Potato leafroll virus* (PLRV) and *Potato virus Y* (PVY) can affect yield substantially, with up to 80% losses, while viruses producing mild or latent symptoms, such as *Potato virus X* (PVX) and *Potato virus S* (PVS), show yield losses of at most 10 to 20% [2]. Host plants can exhibit compatible or incompatible interactions with a virus. In a compatible interaction, potato plants can be either tolerant, accumulating high titres of the virus without symptoms, or sensitive, responding to viral infection with development of disease. In an incompatible interaction, the plants respond to viral infection with a hypersensitive reaction (HR) or an extreme resistance (ER) response. The HR is accompanied by programmed cell death and restricts virus multiplication and spreading. It manifests as necrotic lesions on inoculated leaves and leads to the induction of systemic acquired resistance. The HR is strain-specific and affected by environmental factors (e.g. heat), whereas ER acts against a broad spectrum of virus strains by limiting their accumulation, and only a few or no visible symptoms appear [3]. In potato, HR correlates with the presence of *N* genes, while ER is manifested by *R* genes [4].

Although a relatively large number of virus-resistance genes have been mapped to various chromosomes in potato [5], only a very few of them were isolated and characterised at the DNA sequence level. The first one was the dominant gene *Rx1*, controlling ER to PVX, followed by *Rx2*. Despite their different origins (*S. andigena* and *S. acaule*) and chromosomal locations (XII and V), *Rx1* and *Rx2* share 95% sequence identity [6–8]. At the chromosome XII *Rx1* locus, there are at least three homologues of *Rx1* and the potato cyst nematode-resistance gene *Gpa2*, which is highly similar to *Rx1* [9]. Rx was identified as a protein with a conserved nucleotide binding site and a leucine-rich repeat (NBS-LRR or, shortly, NLR) belonging to the largest class of plant R proteins that can mediate both HR and ER responses [10].

ER against PVY is conferred by *Ry* genes. As early as 1970, five *Ry* genes had been described [11], and three *Ry* genes have already been mapped on potato chromosomes. One of them originates from the wild species *S*. *tuberosum* ssp. *andigena* (*Ry*_*adg*_) and was mapped on chromosome XI [12]. Another one, derived from *S. chacoense* (*Ry*_*chc*_), is located on chromosome IX [13], while the third one, derived from *S. stoloniferum* (*Ry*_*sto*_), was mapped to chromosome XII [14–16]. A gene, designated *Y-1*, co-segregating with *Ry*_*adg*_, was cloned and found to be structurally similar to gene *N* that confers HR to *Tobacco mosaic virus* (TMV) in *Nicotiana* spp. and belongs to the Toll-interleukin-1 receptor (*TIR*)*-type NLR* genes [17]. A *Ry* gene similar to *Y-1* was isolated from the Korean potato cultivar ‘Golden Valley’ and introduced into the ‘Winter Valley’ cultivar which is susceptible to PVY^O^ infection. The transgenic ‘Winter Valley’ showed resistance to PVY^O^ infection [18]. In contrast, leaves of transgenic potato plants expressing *Y-1* under the control of the *CaMV-35S* promoter developed necrotic lesions upon infection with PVY, but no significant resistance was observed, and plants were systemically infected with the virus [17].

Both the environment and evolution modulate viral pathogenesis in plants and *R* genes are in many cases overcome by resistance-breaking strains [19,20]. *Ry_sto_* was introgressed into *S. tuberosum* almost 60 years ago, and various European potato cultivars currently bear *Ry*_*sto*_ [21]. *Ry*_*sto*_ was also introduced into *S. tuberosum* at the Potato Research Centre, Keszthely, Hungary. There has been no indication so far that even the most aggressive PVY strain, NTN, could overcome the ER of *Ry*_*sto*_-bearing potato cultivars [22], including the Hungarian cv. ‘White Lady’ [23]. This phenomenon of unusually durable resistance of *Ry*_*sto*_-bearing potatoes prompted us to map *Ry*_*sto*_ on a fine scale and characterise the *Ry*_*sto*_ locus at the DNA sequence level.

## Materials and methods

### Plant materials and growth conditions

Four hundred fifty-seven genotypes of the tetraploid F1 population described by [23] from a cross between cv. ‘White Lady’ and ‘S440’ were tested for segregation of the *Ry*_*sto*_ gene. The parents and 81 hybrids were obtained from the Potato Research Centre, Keszthely, Hungary, as *in vitro* plants, while the others were grown from seeds. Seed surfaces were disinfected with 20% sodium hypochlorite for 10 min and rinsed with sterile water three times. Seeds were germinated on 1% water-agar Petri plates and placed into 35-ml tubes containing 7 ml RM medium (MS medium without vitamins) [24] containing 2% (w/v) sucrose, solidified with 0.8% agar. Tubes were closed with paper plugs. *In vitro* culturing was performed at 24°C under a light regime of 16 h of light at 75 µmol m^−2^ s^−1^ intensity and 8 h of darkness. Propagation of the plants was carried out *in vitro*.

### Potato transformation

For transformation, the potato cv. ‘Désirée’ was propagated *in vitro* in 500-ml jars in MS medium [24] containing 2% (w/v) sucrose and solidified with 0.8% agar (5 plants/jar). The recombinant vectors from *Escherichia coli* were introduced into *Agrobacterium tumefaciens* strain C58C1 containing pGV2260 [25] by triparental mating [26]. Transgenic ‘Désirée’ lines were generated by leaf transformation according to [27], with 50 µg ml^−1^ kanamycin added to the selection media.

In the case of the potato breeding line ‘S440’, tissue culture-derived sterile microtubers were used for transformation as described by [28] with the exception that the shoots were regenerated and rooted in the presence of 50 µg ml^−1^ kanamycin in the media. Total DNA of putative transgenic plants grown in tissue culture was isolated by the method of [29] and the presence of target genes was verified by PCR using Dream Taq DNA Polymerase (Thermo Fisher Scientific, Waltham, MA, USA) and the gene-specific primers listed in S1 Table with the exception of T2 lines for which a reverse transcription polymerase chain reaction (RT-PCR) was applied as detailed below. PCR-positive transgenic lines from each transformation were propagated *in vitro* and transferred to pots for virus resistance testing.

### Virus resistance testing

For virus tests, four-week-old plants obtained by tissue culture in tubes were transferred into pots and grown further under greenhouse conditions at 20-28°C. After 2-3 weeks, the plants were tested for resistance to PVY^NTN^ by mechanical inoculation. PVY^NTN^ (DSMZ-Deutsche Sammlung von Microorganismen und Zellkulturen GmbH, virus isolate PV-0403) was propagated in *Nicotiana tabacum* cv. Xanthi. Two bottom leaves of potato plants were dusted with carborundum powder, and 100 µl of sap prepared from PVY^NTN^-infected tobacco plant leaves was dropped and dispersed with a micropipette tip onto each leaf. The sap was rubbed into the leaves using a pestle. Non-inoculated upper leaf samples were collected three weeks after inoculation. Detection of the virus was performed by RT-PCR. Total RNA was extracted from leaves according to the method of [30]. RNA was quantified using a NanoDrop spectrophotometer. DNaseI-treated total RNA (1 μg) was reverse-transcribed with RevertAid M-MuLV Reverse Transcriptase and 10xRT Random Primer (Applied Biosystems, Foster City, CA, USA). The cDNAs obtained with the PVY^NTN^ coat protein gene-specific primers (S1 Table) were tested on agarose gel. Hybrids of the F1 population, which appeared initially resistant, were re-tested twice in subsequent experiments. Transgenic ‘Désirée’ and ‘S440’ lines were tested for virus resistance in the same way using three plants per line in each experiment.

### Cloning and sequencing of genetic markers

Cernák et al. [31] developed RAPD markers linked to the *Ry*_*sto*_ gene. The closest RAPD marker amplified from ‘White Lady’ genomic DNA was cloned into the pBluescript SK(+) (Stratagene, La Jolla, CA, USA) and sequenced on an ABI 3100 Genetic Analyser instrument (Biomi Ltd., Gödöllő, Hungary). The SCAR marker ST1 [32] is based on this sequence (S1 Fig.). The YES3-3A marker [33] was also amplified from ‘White Lady’, cloned in pGEM-T Easy (Promega, Madison, WI, USA) and sequenced at Biomi Ltd. (S2 Fig.).

### Bacterial artificial chromosome (BAC) library construction and screening

The BAC library was produced from ‘White Lady’ genomic DNA after partial digestion with *Hind*III in pIndigoBAC-5 at BIO S&T (Montreal, Canada). The total number of clones was 251,160 with an average insert size of 150 Kb. A PCR-based strategy was applied for the identification of BAC clones overlapping the *Ry*_*sto*_ locus. The *Escherichia coli* DH10B carrying the BAC clones in SOC medium [34] supplemented with 12.5 µg ml^−1^ chloramphenicol and 15% glycerol was diluted and divided into 868 x 96 subpools containing approximately 10 individuals each and grown at 37°C in microtiter plates before storage at −80°C. To prepare BAC DNA pools, 124 plates were organised into a composite 3 x 8 grid containing 8 columns and 6 rows. Plasmid DNA was isolated from the 48 pooled samples by the alkaline-lysis method [34] and used as a template for PCR screening of the library. In the case of a positive result, pooled DNA was isolated from the three determined plates and then from the single identified plate. Finally, DNA was prepared and PCR-tested from individual wells. Bacteria of the positive well were plated out and tested individually. PCR primers were designed using Primer3Plus (http://www.bioinformatics.nl/cgi-bin/primer3plus/primer3plus.cgi/) and/or Primer-Blast NCBI (http://www.ncbi.nlm.nih.gov/tools/primer-blast/). After PCR, the products were analysed in an agarose gel.

### Sequencing and bioinformatics analysis of BAC clones

BAC clone DNA was isolated using a Large-Construct Kit (Qiagen, Hilden, Germany). Fragmentation, library production and Illumina MiSeq 2×300 bp sequencing were carried out at the Genomic Medicine and Bioinformatic Core Facility at the University of Debrecen, Hungary. Contig assembly was performed by the A5-miseq pipeline [35]. The raw sequence reads are deposited at the EBI ENA SRA database under the project number PRJEB31027. Publicly available sequence files and other data of potato *S. tuberosum* Group Phureja DM1-3 516R44 originally generated by the Potato Genome Sequencing Consortium [36] were obtained from the Solanaceae Genomics Resource http://solanaceae.plantbiology.msu.edu/pgsc_download.shtml. BAC reads were aligned into the reference genome Phureja using the BWA-MEM program [37]. Multiple sequence alignments were carried out by BLASTn (https://blast.ncbi.nlm.nih.gov/) and Clustal Omega (https://www.ebi.ac.uk/Tools/msa/clustalo/). The predictions of open reading frames and exon-intron boundaries were based on the GENSCAN webserver at MIT (http://genes.mit.edu/GENSCAN.html).

### Cloning of candidate genes

Cloning of candidate genes was based on PCR using the primers listed in S1 Table. Long-range PCR amplifications were performed using Phusion High-Fidelity DNA Polymerase (Thermo Fisher Scientific, Waltham, MA, USA). The *DisRes* fragment was cloned into the *Sma*I site of the binary vector pBin19, providing kanamycin resistance in transgenic plants [38]. In all other cases, the PCR primers were extended at the ends with recognition sites for *Bam*HI and inserted into the *Bam*HI site of pBin19. The recombinant vectors were transformed into the *Escherichia coli* strain DH5α [39].

## Results

### Fine-scale genetic mapping of the *Ry*_*sto*_ gene

According to the previous result obtained by [31], the tetraploid potato cultivar ‘White Lady’ carries the *Ry*_*sto*_ gene in simplex form. In this study, among the 457 tested F1 genotypes derived from a cross between ‘White Lady’ and ‘S440’, 220 resistant plants and 237 plants sensitive to PVY^NTN^ infection were identified. The segregation ratio of 1:1 confirmed the presence of a single, dominant gene for extreme resistance to PVY^NTN^ in simplex state in the tetraploid parental variety ‘White Lady’.

Molecular markers co-segregating with *Ry*_*sto*_, e.g., STM0003, YES3-3A, Cat-in2 and ST1, were identified earlier [16,31-33,40]. The closest markers to *Ry*_*sto*_ in the genetic map of ‘White Lady’ were STM0003 and ST1 in a distance of 2.95 and 0.53 cM, respectively [31]. Since STM0003 seemed to be too far from *Ry*_*sto*_ to start chromosome walking towards the resistance gene a fine mapping was carried out by testing the co-inheritance of the above listed four molecular markers with the virus resistant/sensitive phenotype in 400 genotypes derived from the ‘White Lady’ x ‘S440’ cross. Seven recombinants between STM0003 and Cat-in2 were identified. Thus, the genetic distance between the two markers was estimated to be 1.75 cM. STM0003 at the proximal site and Cat-in2 at the distal site surrounded the *Ry*_*sto*_ gene. Based on the recombinant events detected in the seven recombinants (Fig. 1), the order of the four markers, STM0003, YES3-3A, ST1 and Cat-in2, was established as shown in Fig. 2.

**Fig. 1.**
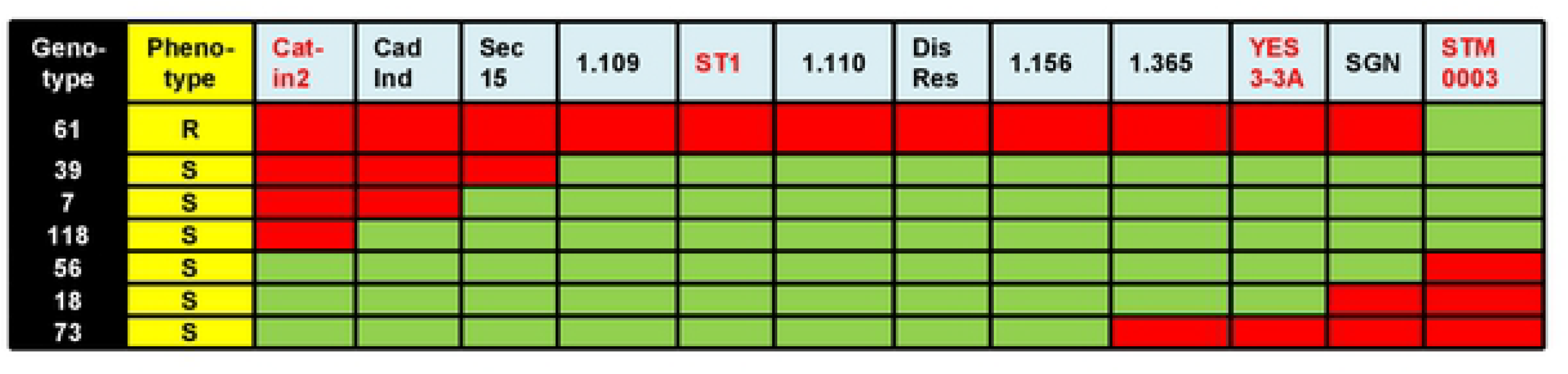
Colourmap of the *Ry*_*sto*_ region. R, resistant; S, sensitive. The presence of markers representing the resistant parent ‘White Lady’ are indicated by red boxes, while the presence of markers representing the sensitive parent ‘S440’ are indicated by green boxes in seven genotypes of the F1 population. Published markers are red-coloured.

**Fig. 2.**
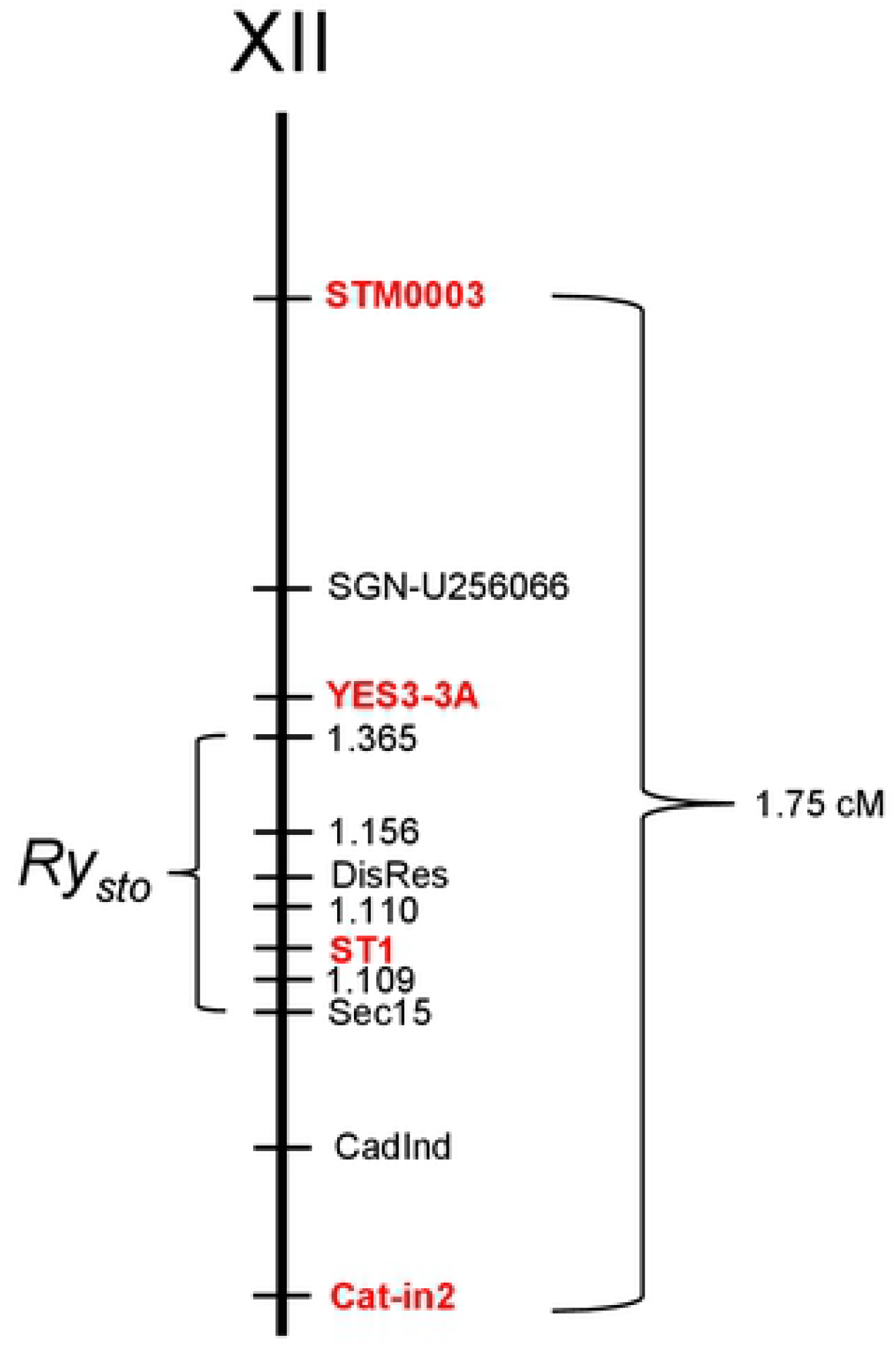
Position of the *Ry*_*sto*_ locus on chromosome XII. The genetic distance in cM is shown on the right. The map distances for STM0003, SGN-U256066, YES3-3A, 1.365, Sec15, CadInd and Cat-in2 were calculated from recombination frequencies between DNA markers and resistance loci, whereas localisation of 1.156, DisRes, 1.110, ST1, and 1.109 were based on the Phureja genome sequence. Published markers are red-coloured.

### Identification of supercontigs carrying the *Ry*_*sto*_ locus

The availability of the *S. tuberosum* Group Phureja genome sequence (Potato Genome Sequencing Consortium 2011) provided the possibility for physical mapping of the *Ry*_*sto*_ locus. Browsing the PGSC database, we found the STM0003 marker on the supercontig PGSC0003DMB000000114. To position additional markers of the *Ry*_*sto*_ locus on supercontigs, the ST1 and YES3-3A PCR fragments of ‘White Lady’ were cloned and sequenced (S1,2 Fig.). DNA sequence comparison localised YES3-3A to the same supercontig as STM0003, while ST1 identified PGSC0003DMB000000034.

The tomato (*S. lycopersicum*) genome sequence was also available in the database (The Tomato Genome Consortium, 2012), and we found that a large number of markers were localised to the tomato region orthologous to the identified potato supercontigs. Based on these tomato markers, 16 primer pairs were synthesised and tested; however, only one of them, designated SGN-U256066, showed polymorphism between ‘White Lady’ and ‘S440’. This marker was located between STM0003 and YES3A (Fig. 1 and 2).

### Isolation of BAC clones overlapping the *Ry*_*sto*_ locus

A BAC library was constructed from the genomic DNA of ‘White Lady’. Physical mapping started with screening the BAC library using the ST1 marker by which five positive BACs were identified (Fig. 3). In a subsequent experiment, YES3-3A identified one BAC clone (Fig. 3). The ends of the six positive BACs were sequenced. Sequences were mapped to the *S. tuberosum* Group Phureja genome sequence. The comparison localised the *Ry*_*sto*_ locus to the 57-59 Mb segment of chromosome XII in Phureja. The obtained sequences were used for new marker development. A screen for markers in the intergenic regions successfully identified a polymorphic marker designated 1.365. Genetic mapping of 1.365 reduced the size of the *Ry*_*sto*_ locus and localised *Ry*_*sto*_ between Cat-in2 and 1.365 (Fig. 1, 2 and 3). To close the genetic window, further PCR primers were designed based on the Phureja genome sequence and tested for polymorphism between ‘White Lady’ and ‘S440’. Three new markers were found in this way: CadInd, Sec15 and DisRes. Testing the six lines bearing recombination between Cat-in2 and 1.365 with the new markers, the *Ry*_*sto*_ locus could be narrowed to the Sec15-1.365 fragment between 58 and 59 Mb (Figs. 1, 2 and 3). BAC walking was continued with Sec15, DisRes and 1.365 markers and resulted in the isolation of five new clones, two by DisRes and three by 1.365 (Fig. 3). No BACs were isolated by Sec15 in the 48 pooled samples tested. To close the gap between the isolated BAC clones, an attempt was made to identify new polymorphic markers based on the end sequences of BAC inserts. This attempt resulted in the identification of markers 1.109, 1.110 and 1.156 (Figs. 1 and 2). In comparison to the Phureja genome sequence, isolation of one BAC clone by 1.109 and another clone by 1.110 closed the gap between the BACs overlapping the *Ry*_*sto*_ locus (Fig. 3).

**Fig. 3.**
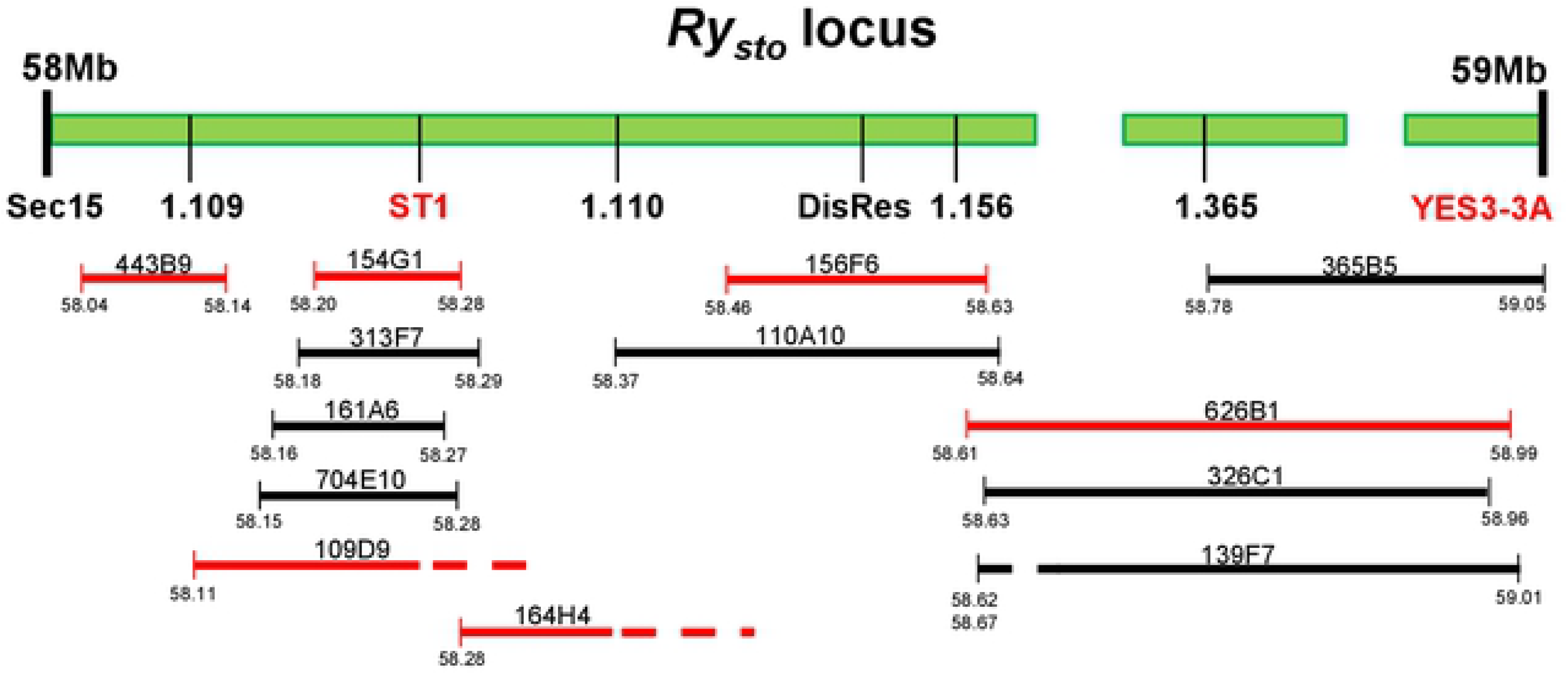
Physical map of the *Ry*_*sto*_ locus with the overlapping BAC clones. The *Ry*_*sto*_ locus in the *S. tuberosum* cv. ‘White Lady’ corresponds to the 58-59-Mb region on chromosome XII in the genome-sequenced *S. tuberosum* Group Phureja DM1-3 516R44. This region of Phureja has sequence gaps probably due to its highly repetitive nature. Markers used for genetic mapping and isolation of BAC clones are indicated under the upper line illustrating the chromosomal fragment. BAC clones are represented by the horizontal lines with their names on them. The published markers and the fully sequenced BAC clones are highlighted in red. The numbers under the lines indicate the position of the BAC ends on the corresponding Phureja genome sequence. The position of BAC ends indicated by dashed lines could not be defined precisely because these segments possess a large number of repeated sequences.

### Selection of candidate genes by DNA sequence similarity

Six presumably overlapping BAC clones, namely, 443B9, 109D9, 154G1, 164H4, 156F6 and 626B1 (Fig. 3), were sequenced. The reads were *de novo* assembled both pooled and individually by the A5-miseq pipeline. The pooled BAC reads gave 49 contigs, which were greater than 3 Kb. The sum length of these contigs was 0.97 Mb out of which 0.38 Mb could be aligned to the ~1 Mb region of Phureja corresponding to the *Ry*_*sto*_ locus in ‘White Lady’ (S3 Fig.), while 243 Kb mostly repetitive sequences were mapped to other regions of the Phureja genome. Interestingly, 313 Kb did not map at all to the Phureja reference genome.

Sequences of 13 out of the 16 *Ry*_*sto*_ locus-specific primers co-segregating with the *Ry*_*sto*_-provided extreme PVY^NTN^ resistance (Fig. 1) possessed 100% identity with the corresponding BAC clone sequences and only 1-2 bp difference was detected in the case of the other three primers (S5 Fig.) indicating that all six BAC clones were originated from the *Ry*_sto_-bearing chromosome.

The end regions between the six sequenced BAC clones were examined pair-wise. As expected, the 443B9, 109D9 and 154G1 BAC clones showed 100% identity in the overlapping regions. Unfortunately, the overlapping region between the 154G1 and 164H4 BAC clones was restricted only to 8 bp including a *Hind*III site, which served as the cloning site during generation of the BAC library. In order to demonstrate the adjacent position of the two BAC clones a primer pair complementary to the corresponding ends of 154G1 and 164H4 was designed and used in PCR with ‘White Lady’ genomic DNA as a template. The reaction resulted in an approximately 0.45 kb PCR product as it was visualised on an agarose gel. The PCR product was cloned and Sanger-sequenced. Clones representing all four chromosomes were obtained with no additional sequences compared to the 8-bp overlap between 154G1 and 164G1 (S4 Fig.) indicating that 154G1 and 164H4 represent a continuous fragment of the *Ry*_*sto*_ locus. On the other side, the end of the 164H4 BAC clone showed 100% identity with the overlapping 156F6 contigs. The 156F6 BAC clone, however, did not overlap with the 626B1 contigs. Although, the BWA-MEM alignment showed high similarity between the two BAC ends (S3 Fig.), the NCBI BLAST revealed that the similarity was due to the presence of *NLR* homologous sequences of truncated genes on the tested scaffolds, while their very ends were different.

Based on DNA sequence comparison to the annotated Phureja genome and searches in the NCBI database, eight genes were assigned to the *NLR* family. Seven *NLR* genes were located on the BAC clone 156F6 and one on 626B1. *NLR* genes can be divided into two subclasses: one includes genes whose proteins contain a coiled-coil (CC) motif at their N-terminus, and the other includes genes whose proteins resemble the Toll-interleukin receptor (TIR) domain at the N-terminus [10]. One out of the eight *NLR* genes that we identified was a *CC-type* disease resistance gene encoding a protein 97% identical to the predicted disease resistance RPP8-like protein 2 of potato (S6 Fig.). Since this gene carried the DisRes marker we kept the name *DisRes* for the *RPP8*-like gene located on BAC clone 156F6. The other seven putative resistance genes belonged to the *TIR-type NLR* genes. These included four genes encoding proteins similar to the phloem protein A5-like. Nevertheless, one out of the four genes that we identified (*Phloem3*) may not be functional because it encodes an N-terminal-truncated protein and involves four stop codons. The other three genes, *Phloem 1, 2* and *4*, encoded 77-87% identical proteins (S7 Fig.). The other three *TIR-type NLR* genes were similar to the *N*-like *TMV resistance* gene isolated from tobacco (S8 Fig.). Thus, these genes were designated *TMV1*, *TMV2* and *TMV3*. The predicted proteins TMV2 and TMV3 possessed 80% identity at the amino acid sequence level. *TMV1*, however, similar to *Phloem3*, may not be functional because it encodes an N-terminal-truncated protein (S8 Fig.).

Vidal et al. [17] cloned and characterised *Y-1* that co-segregated with *Ry*_*adg*_, a gene for ER to PVY on chromosome XI and found it also structurally similar to the *N*-like *TMV resistance* gene isolated from tobacco. Therefore, we tested the similarities between *Y-1* and the *TMV*s isolated from ‘White Lady’. The highest identity, 42%, was detected between Y-1 and TMV2 (S9 Fig.). Recently, Grech-Baran et al. [42] published the genomic sequence of a *TIR-NLR* immune receptor identified as *Ry*_*sto*_ in a dihaploid clone of the cultivar “Alicja”, which has PVY resistance also from *S. stoloniferum* in its ancestry. Comparison of the two genomic sequences from the putative start and stop codons of the genes revealed 95% identity between *TMV2* and *Ry*_*sto*_ (S10 Fig.).

### Cloning and functional testing of *Ry*_*sto*_ candidates

The putative resistance genes were subcloned from the BAC clone 156F6 into the binary vector pBin19 for *Agrobacterium*-mediated transformation of the PVY susceptible potato cultivars ‘Désirée’ and ‘S440’. Six genes of interest with 0.4-1.9 kb untranslated 5’ regions were PCR-amplified with a high fidelity DNA polymerase and inserted into pBin19. *TMV2, TMV3* and *DisRes* were cloned separately, resulting in the constructs T2, T3 and DR, while *Phloem* genes were cloned in pairs, resulting in the constructs P1-2 and P3-4 (Fig 4).

**Fig. 4.**
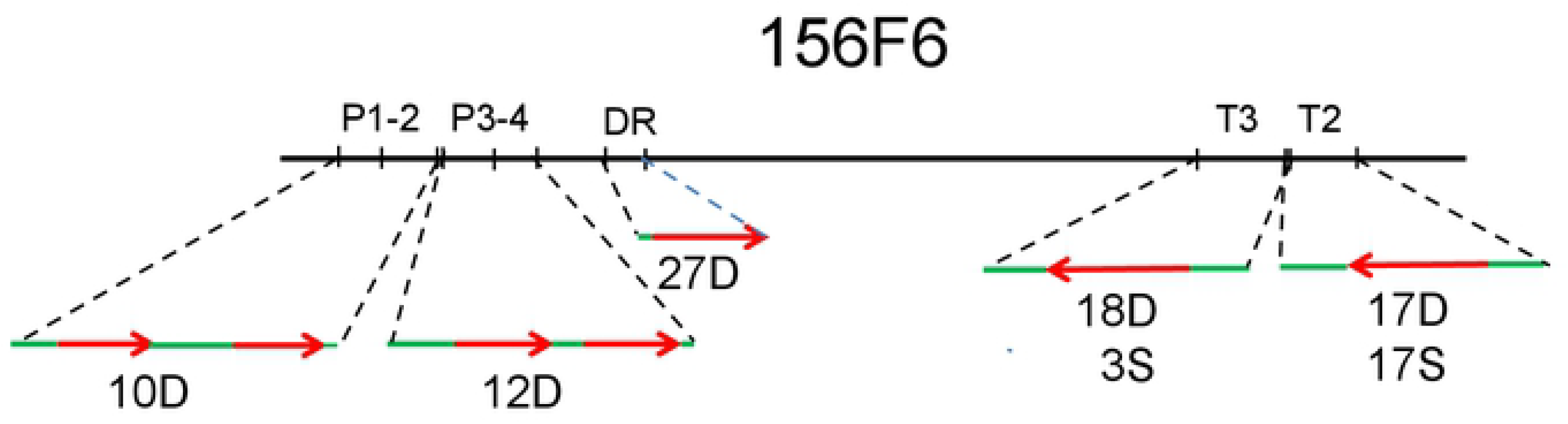
Schematic representation of the putative *NLR* resistance genes on the BAC clone 156F6 and the constructs used for transformation with the number of transgenic lines tested for virus resistance. Red arrows indicate the coding regions of the genes with introns and their direction of transcription. Green lines represent the non-coding regions. The genomic fragments were cloned in the binary vector pBIN19 and transformed into the PVY-sensitive potato cultivar ‘Désirée’ and ‘S440’. Symbols: P, *phloem protein-coding* gene; DR *disease resistance* gene; T, *TMV resistance* gene; D, ‘Désirée’; S, ‘S440’.

Twenty to forty ‘Désirée’ leaves per construct were transformed and 80-100 plants regenerated and rooted on selective media indicating that the transformations were efficient. Twelve independent putative transgenic plants (i.e., plants regenerated from different leaves) derived from the transformation with P1-2 and P3-4 each, and 20, 24 and 30 plants from the transformation with *TMV2*, *TMV3* and *DisRes*, respectively, were isolated. To test for the presence of ‘White Lady’-derived genes in the putative transgenic plants gene-specific primers were developed (Table 1 and Fig. 5). DNA was isolated from *in vitro* grown P1-2, P3-4, DR and T3 plants and the gene-specific primers were used for PCR amplification of *NLR* fragments. In this way, 10 P1-2, 12 P3-4, 27 DR and 18 T3 lines were selected. In the case of *TMV2* no primer pair distinguishing between the corresponding ‘White Lady’ and ‘Désirée’ gene could be identified. Nevertheless, with a primer pair designed for *TMV2* cDNA (Table 1) a difference in expression level of the gene was detected (Fig. 6). Thus in the case of T2 lines, RNA was isolated from the leaves, and RT-PCR was used to identify the 17 ‘Désirée’ transgenic lines expressing *TMV2.* Attempts were made to detect the expression of the other transgenes as well, however, due to the high level of homology (S6,7 Fig.) no cultivar-specific primers for the *phloem protein-like* and *DisRes* transcripts could be designed, while in the case of *TMV3* the level of expression was very low even in ‘White Lady’.

**Table 1.**
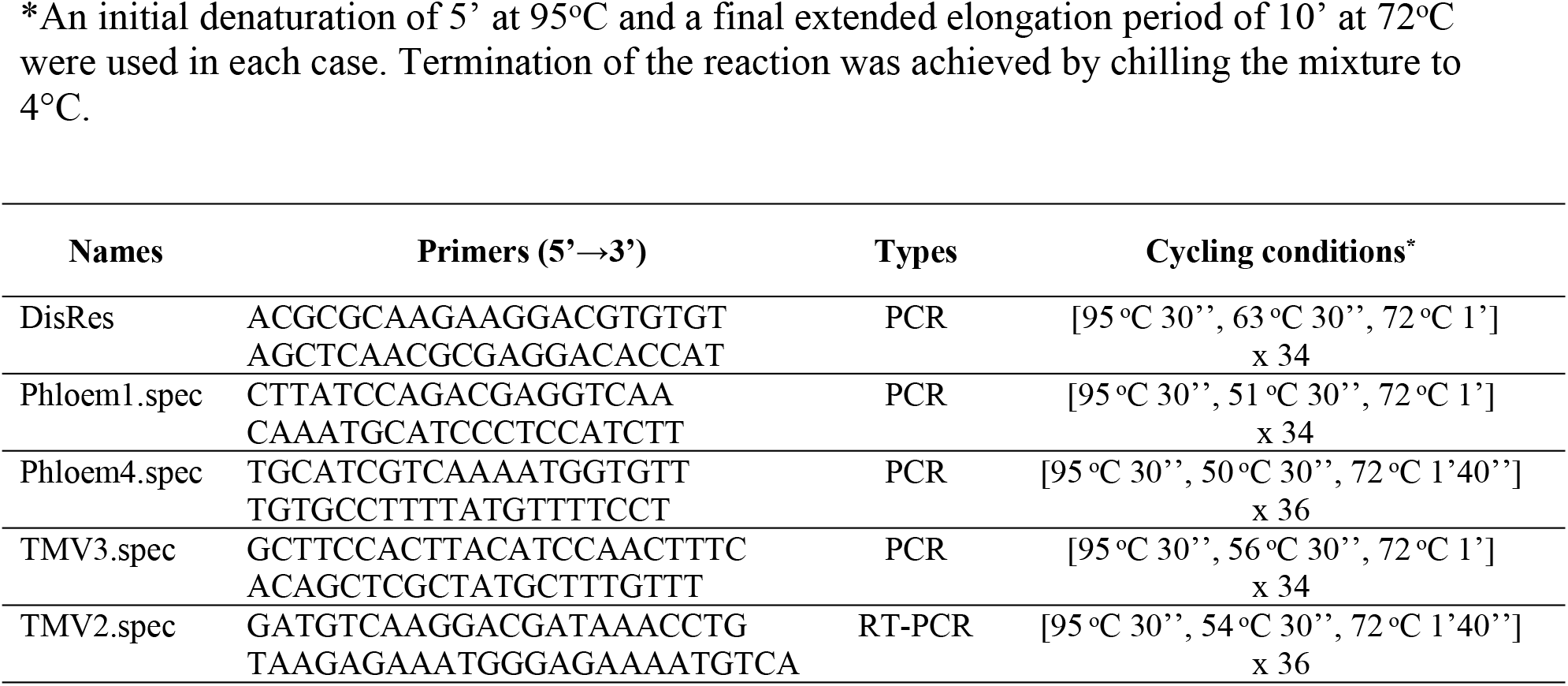
*NLR* gene-specific markers of the *Ry*_*sto*_ locus *An initial denaturation of 5’ at 95°C and a final extended elongation period of 10’ at 72°C were used in each case. Termination of the reaction was achieved by chilling the mixture to 4°C.

**Fig. 5.**
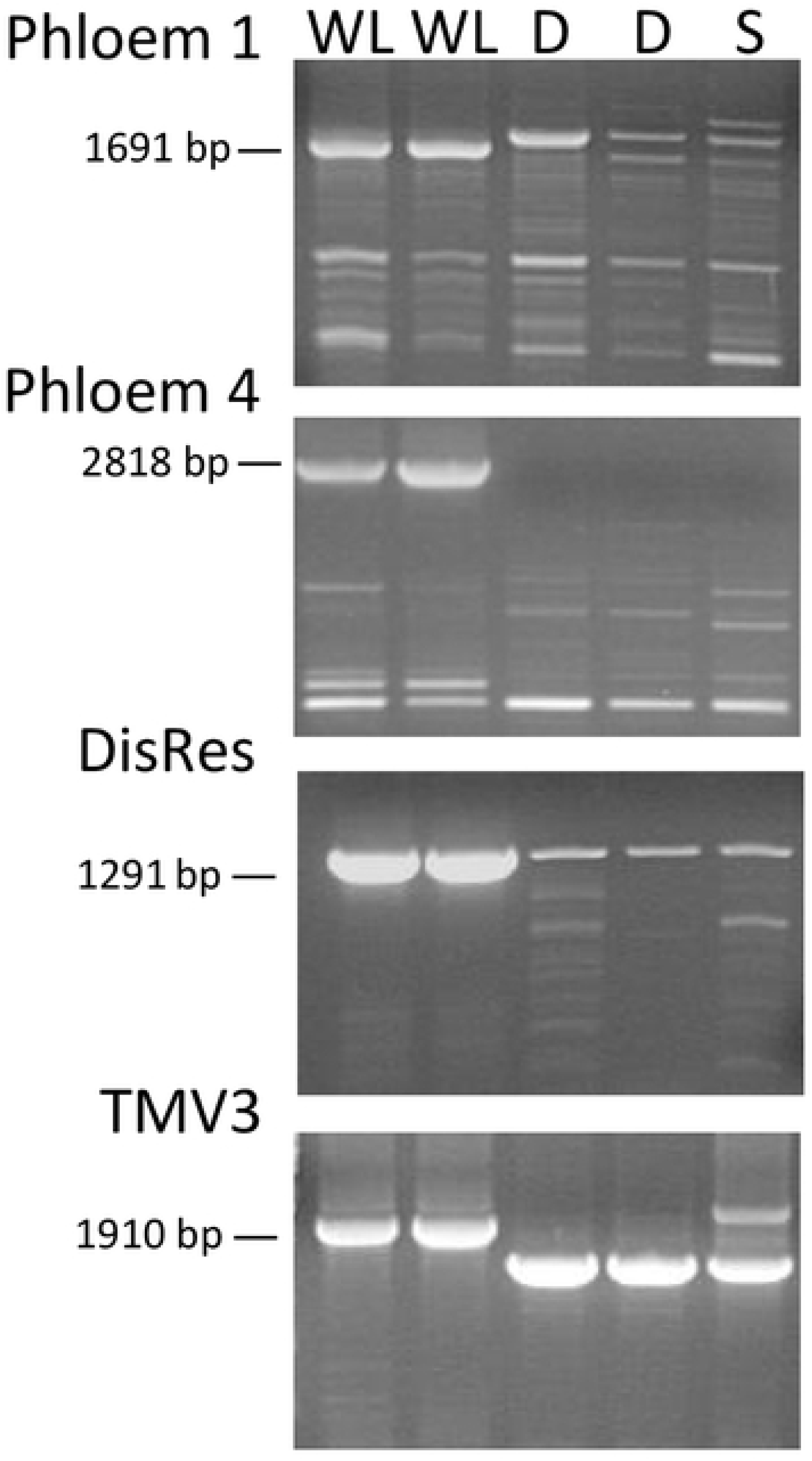
*NLR*-specific markers tightly linked to the *Ry*_*sto*_ gene as detected on agarose gel. WL, ‘White Lady’; D, ‘Désirée’; S, ‘S440’. PCR fragments were generated from genomic DNA with the primer pairs presented in Table 1.

**Fig. 6.**
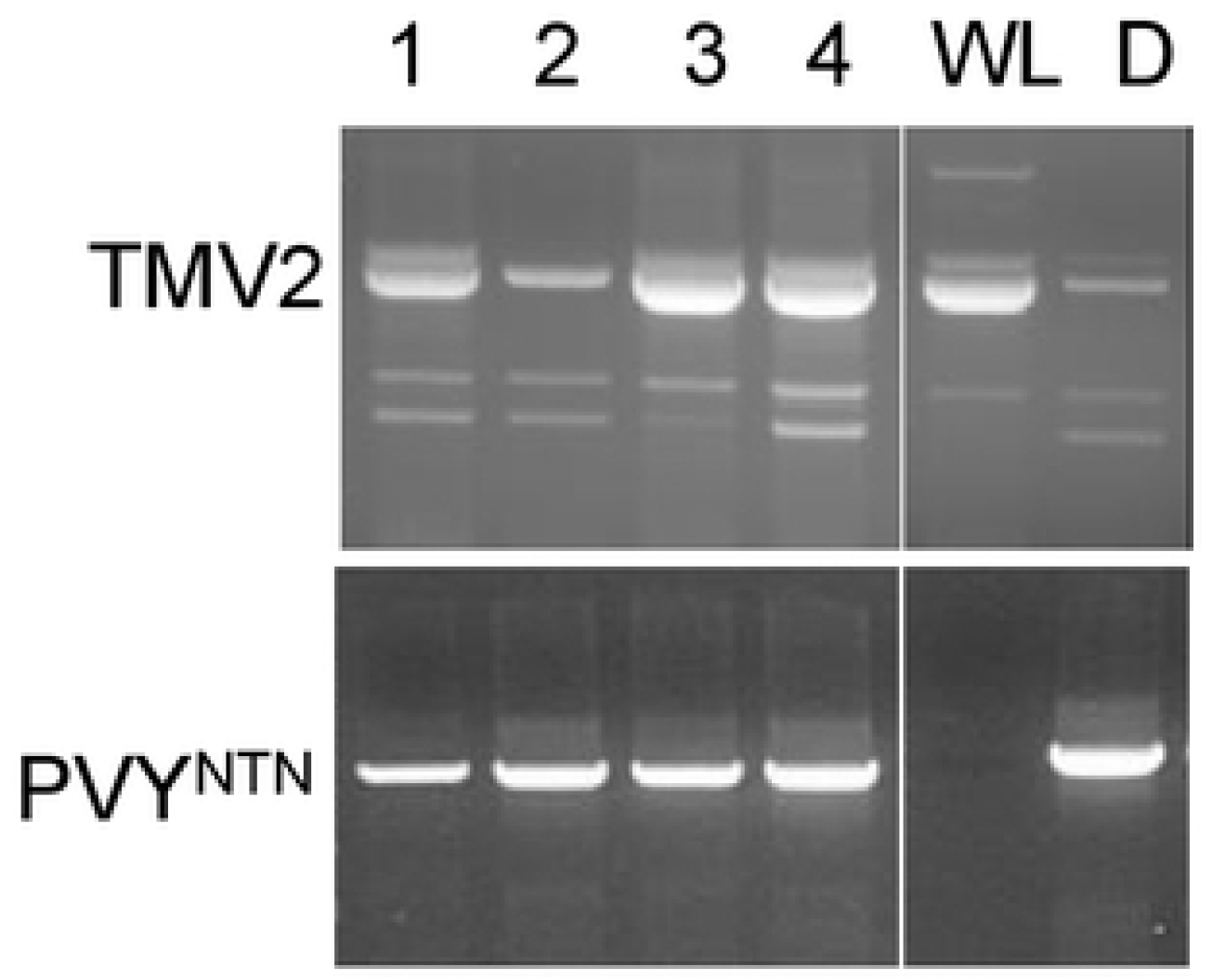
Detection of *TMV2* expression and PVY^NTN^ in ‘Désirée’ plants transformed with the T2 construct. RNA was isolated from the upper leaves of plants three weeks after viral infection of bottom leaves. RT-PCR fragments were generated with *TMV2*- and *PVY coat protein* gene-specific primers and separated on agarose gels. Lines 1, 3 and 4 were considered positive, while line 2 was considered negative for the expression of the transgene. Symbols: WL, ‘White Lady’; D, ‘Désirée’.

To test the virus sensitivity of the transgenic plants the selected lines were propagated *in vitro*, planted in pots and inoculated with PVY^NTN^ under greenhouse conditions. Unfortunately, none of the transgenic ‘Désirée’ lines proved to be virus resistant in repeated experiments. The example of a few TMV2 transgenic lines is shown in Fig. 6.

An attempt was made to introduce *TMV2* and *TMV3* not only into ‘Désirée’ but also into ‘S440’. Nevertheless, leaf transformation using the same method used for ‘Désirée’ failed for ‘S440’. Therefore, sterile ‘S440’ microtubers were obtained from *in vitro* plants and transformed with T2 and T3 constructs. Seventeen T2 and three T3 transgenic lines were isolated and subjected to virus testing, but no significant resistance to PVY^NTN^ was observed.

## Discussion

The *Ry*_*sto*_ gene from *S. stoloniferum* was originally introgressed into the widely used breeding clone MPI 61.303/34. *Ry*_*sto*_-based virus resistance has proven quite durable and is used in breeding programmes throughout the world. For example, this gene provides resistance against several viruses in the cultivars ‘Bzura’, ‘Forelle’, ‘Pirola’, and ‘White Lady’ [43], the last of which is the object of this study.

Our result supported the previous finding [16] that the *Ry*_*sto*_ gene in ‘White Lady’ is located on chromosome XII and linked to STM0003 and YES3-3A, the markers widely used for marker-assisted selection in breeding programmes [44,45]. Because *Ry-f*_*sto*_ mapped by Flis et al. [14] and Song et al. [15] is also linked to STM0003 and YES3-3A, there is a possibility that the *Ry*_*sto*_ gene in ‘White Lady’ and *Ry-f*_*sto*_ have the same source.

In addition to the abovementioned publications, some other studies also supported the location of *Ry*_*sto*_ on the distal end of chromosome XII [21,46,47]. To the best of our knowledge, however, the genetic map presented in Fig. 2 is the most detailed map of the *Ry*_*sto*_ locus published so far. Sequencing the PCR fragments of ‘White Lady’ generated by ST1 and YES3-3A primers identified two supercontigs as a putative region surrounding *Ry*_*sto*_ and resulted in the isolation of a new marker, SGN-U256066, located between STM0003 and YES3-3A. Combining the genetic map with genomic sequence data of the *S. tuberosum* Group Phureja, the size of the *Ry*_*sto*_ locus was narrowed down to approximately 1 Mb. BAC clones of ‘White Lady’ overlapping the *Ry*_*sto*_ locus were isolated by chromosome walking. Markers 1.365, 1.156, 1.110 and 1.109 were designed to the BAC clone ends and used in walking. For the identification of two additional markers, Sec15 and DisRes, the intron targeting (IT) method was applied. This method is based on the observation that intron sequences are generally less conserved than exons, and they display polymorphism due to length and/or nucleotide variation in their alleles. Effective strategies for exploiting this information and generating IT markers have already been developed and successfully applied for many plant species, including potato [31,48].

By annotating the *Ry*_*sto*_ locus sequence, one putative *CC-type* and seven *TIR-type NLR* genes were identified, of which five appeared to encode full-length proteins. A genome-wide genetic mapping of *NLR* disease resistance loci in the diploid potato clone RH89-039-16 (*S. tuberosum* ssp. *tuberosum*) resulted in the detection of 738 partial- and full-length resistance gene homologues [46]. A very similar result was obtained by resistance gene enrichment sequencing (RenSeq), which detected 755 *NLR*s in the sequenced *S. tuberosum* genome [49]. Nevertheless, the function of most of these genes is unknown. In our case, however, the *CC-type NLR* was similar to *RPP8*, a gene of *Arabidopsis thaliana* providing resistance to *Peronospora parasitica* [50]. In addition, three genes encoding full-length proteins similar to phloem protein A5-like were identified. It was shown earlier that many phloem proteins have roles in wound and defence responses [51]. Two genes encoding full-length proteins were similar to TMV resistance proteins in tobacco [52] and Y-1 characterising an ER with *S. andigena* origin in potato [17]. One of these genes, *TMV2*, showed 95% identity with the recently isolated *Ry*_*sto*_ gene of the dihalpoid clone dH ‘Alicja’ at genomic DNA sequence level [42].

Implication of the *CC-type NLR* gene *DisRes* and the five *TIR-type NLR* genes, *Phloem 1, 2, 4* and *TMV2, 3*, in ER was tested by introducing the genes into the PVY-sensitive potato cultivar ‘Désirée’ and expressing the genes using their own putative promoter. The presence of the *NLR* genes derived from ‘White Lady’ was demonstrated in transgenic lines selected for virus resistance testing; however, the expression of *TMV2* only could be tested in transgenic plants due to the absence of *phloem protein-like* and *DisRes* cDNA-specific primers and the very low level of expression of *TMV3* in ‘White Lady’. None of the transgenic lines, however, became PVY^NTN^ resistant. A similar result was obtained by [17] while testing *Y-1*, the gene that co-segregated with the ER gene *Ry*_*adg*_. The transgenic potato plants of line v2-134 expressing *Y-1* under the control of *CaMV-35S* promoter developed necrotic lesions upon infection with PVY^O^, but no significant resistance was observed, and plants were systemically infected with the virus. Thus, it was hypothesised that the function of Y-1 is merely to cause a PVY-specific cell death response.

In contrast, introduction and expression of a *TIR-NLR* gene derived from ‘Alicja’ and mapped to the same 58-59-Mbp region of chromosome XII as the *TIR-NLR* genes we identified transferred ER to two sensitive potato cultivars, ‘Russet Burbank’ and ‘Maris Piper’. This gene, which was identified as *Ry*_*sto*_ in ‘Alicja’ [42], is 95% identical with *TMV2* in ‘White Lady’ at genomic sequence level. Even the primer pair used to clone *Ry*_*sto*_ from ‘Alicja’ may be suitable for cloning *TMV2* from ‘White Lady’ (S10 Fig.). Therefore, it is highly probable that *TMV2* is the *Ry*_*sto*_ gene in ‘White Lady’. Nevertheless, expression of *TMV2* did not confer ER neither to transgenic ‘Désirée’ nor to ‘S440’ lines. Although a high-fidelity DNA polymerase was used to amplify *TMV2* from the ‘White Lady’ genome a mismatch caused by the enzyme could result in the loss of function of the gene. An alternative explanation might be the high frequency of chimera formation generated by *Agrobacterium*-mediated transformation and regeneration from calluses [53]. Since a slightly different transformation technique was used by Grech-Baran et al. [42] and us, and the frequency of chimera formation may be different in various potato cultivars, these factors could result in a difference in the frequency of chimera formation. Since chimeras have untransformed cells the viruses could spread and multiply in these cells masking the resistant phenotype of the transgenic cells when testing the plants for the presence of the virus by RT-PCR. It should be noted, however, that based on DNA sequence analysis there is a gap between the BAC clones 156F6 and 626B1 in our map. Thus, the presence of a functional gene corresponding to *Ry*_*sto*_ and located in the uncovered region cannot be excluded either.

Marker-assisted selection (MAS) has already been routinely employed in crop breeding programmes to accelerate cultivar development. With selection for virus resistance in the juvenile phase and parental selection prior to crossing, breeding time and costs can be reduced. In this work, we discovered five new markers very tightly linked to *Ry*_*sto*_, which showed polymorphism between the *Ry*_*sto*_ bearing potato cv. ‘White Lady’ and the virus sensitive ‘Désirée’ and ‘S440’ (Fig. 5 and 6). In addition to the markers identified earlier, these markers can serve potato breeding efforts for extreme virus resistance.

## Supporting information

**File S1** S1 Table. List of primers. (pdf)

**File S2 Supplementary figures. S1 Fig.** DNA sequence of ST1; **S2 Fig.** DNA sequence of YES3-3A; **S3 Fig.** Alignment of BAC clone sequences to Phureja sequence; **S4 Fig.** End sequences of the BAC clones 154G1 and 164H4; **S5 Fig.** DNA sequence identity of the *Ry*_*sto*_ region specific primers with the corresponding BAC clone sequences; **S6 Fig.** Amino acid alignment of DisRes with RPP8-like protein 2; **S7 Fig.** Amino acid alignment of phloem proteins; S8 Fig. Amino acid alignment of TMV resistance proteins; **S9 Fig.** Amino acid alignment of TMV2 with Y-1; **S10 Fig.** DNA sequence alignment of the *TMV2* genomic region of ‘White Lady’ and the *Ry*_*sto*_ region of dH ‘Alicja’. (pdf)

## Acknowledgements

The authors are grateful to M. Kiss for the excellent technical help in propagation and transformation of potato plants.

## References

1. Jones JD, Witek K, Verweij W, Jupe F, Cooke D, Dorling S, et al. Elevating crop disease resistance with cloned genes. Philos Trans R Soc Lond B Biol Sci. 2014;369: 20130087.

2. Palukaitis P. Resistance to viruses of potato and their vectors. Plant Pathol J. 2012;28: 248–258.

3. Muthamilarasan M, Prasad M. Plant innate immunity: an updated insight into defense mechanism. J Biosci. 2013;38: 433–449.

4. Kopp A, Kondrák M, Bánfalvi Z. Molecular mechanisms of resistance to *Potato virus X* and *Y* in potato. Acta Physiol Entomol Hung. 2015;50: 151–160.

5. Solomon-Blackburn RM, Barker H. A review of host major-gene resistance to potato viruses X, Y, A and V in potato: genes, genetics and mapped locations. Heredity (Edinb). 2001;86: 8–16.

6. Ritter E, Debener T, Barone A, Salamini F, Gebhardt C. RFLP mapping on potato chromosomes of two genes controlling extreme resistance to potato virus X (PVX). Mol Gen Genet. 1991;227: 81–85.

7. Bendahmane A, Kanyuka K, Baulcombe DC. The *Rx* gene from potato controls separate virus resistance and cell death responses. Plant Cell. 1999;11: 781–791.

8. Bendahmane A, Querci M, Kanyuka K, Baulcombe DC. *Agrobacterium* transient expression system as a tool for the isolation of disease resistance genes: application to the *Rx2* locus in potato. Plant J. 2000;21: 73–81.

9. van der Vossen EA, van der Voort JN, Kanyuka K, Bendahmane A, Sandbrink H, Baulcombe DC, et al. Homologues of a single resistance-gene cluster in potato confer resistance to distinct pathogens: a virus and a nematode. Plant J. 2000;23: 567–576.

10. Sukarta OC, Slootweg EJ, Goverse A. Structure-informed insights for NLR functioning in plant immunity. Semin Cell Dev Biol. 2016;56: 134–149.

11. Cockerham G. Genetical studies on resistance to Potato viruses X and Y. Heredity. 1970; 25: 309–348.

12. Hamalainen JH, Watanabe KN, Valkonen JPT, Arihara A, Plaisted RL, Pehu E, et al. Mapping and marker-assisted selection for a gene for extreme resistance to potato virus Y. Theor Appl Genet. 1997;94: 192–197.

13. Hosaka K, Hosaka Y, Mori M, Maida T, Matsunaga H. Detection of a simplex RAPD marker linked to resistance to potato virus Y in a tetraploid potato. Am J Pot Res. 2001;78: 191–196.

14. Flis B, Hennig J, Strzelczyk-Zyta DS, Gebhardt C, Marczewski W. The *Ry-f*_*sto*_ gene from *Solanum stoloniferum* for extreme resistant to Potato virus Y maps to potato chromosome XII and is diagnosed by PCR marker GP122_718_ in PVY resistant potato cultivars. Mol Breeding. 2005;15: 95–101.

15. Song Y-S, Hepting L, Schweizer G, Hartl L, Wenzel G, Schwarzfischer A. Mapping of extreme resistance to PVY (*Ry*_*(sto)*_) on chromosome XII using anther-culture-derived primary dihaploid potato lines. Theor Appl Genet. 2005;111: 879–887.

16. Cernák I, Decsi K, Nagy S, Wolf I, Polgár Z, Gulyás G, et al. Development of a locus-specific marker and localization of the *Ry*_*sto*_ gene based on linkage to a catalase gene on chromosome XII in the tetraploid potato genome. Breeding Sci. 2008;58: 309–314.

17. Vidal S, Cabrera H, Andersson RA, Fredriksson A, Valkonen JP. Potato gene *Y-1* is an *N* gene homolog that confers cell death upon infection with *potato virus Y*. Mol Plant Microbe Interact. 2002;15: 717–727.

18. Lee C, Park J, Hwang I, Park Y, Cheong H. Expression of *G-Ry* derived from the potato (*Solanum tuberosum* L.) increases PVY^(O)^ resistance. J Agric Food Chem. 2010;58: 7245–7251.

19. Fraile A, García-Arenal F. Environment and evolution modulate plant virus pathogenesis. Cur Opin Virol. 2016;17: 50–56.

20. Boualem A, Dogimont C, Bendahmane A. The battle for survival between viruses and their host plants. Curr Opin Virol. 2016;17: 32–38.

21. Valkonen, JPT, Wiegmann K, Hämäläinen JH, Marczewski W, Watanabe KN. Evidence for utility of the same PCR-based markers for selection of extreme resistance to Potato virus Y controlled by *Ry*_*sto*_ of *Solanum stoloniferum* derived from different sources. Ann Appl Biol. 2008;152: 121–130.

22. Davie K, Holmes R, Pickup J, Lacomme C. Dynamics of PVY strains in field grown potato: Impact of strain competition and ability to overcome host resistance mechanisms. Virus Res. 2017;241: 95–104.

23. Horváth S, Wolf I, Polgár Z. Results and importance of resistance breeding against viruses in Hungary. In Abstracts of International Symposium, Breeding Research on Potatoes, Rostock Germany, 23-26 June, 1998, Book of Proceedings; 1998; pp. 75–80.

24. Murashige T, Skoog F. A revised medium for rapid growth and bioassays with tobacco tissue cultures. Physiol Plant. 1962;15: 473–497.

25. Deblaere R, Bytebier B, De Greve H, Deboeck F, Schell J, van Montagu M, et al. Efficient octopine Ti plasmid-derived vectors for *Agrobacterium*-mediated gene transfer to plants. Nucl Acids Res. 1985;13: 4777–4788.

26. Ditta GS, Stanfield D, Corbin D, Helinski DR. Broad host-range DNA cloning system for gram-negative bacteria: construction of a gene bank of *Rhizobium meliloti*. Proc Natl Acad Sci USA. 1980;77: 7347–7351.

27. Dietze J, Blau A, Willmitzer L. *Agrobacterium*-mediated transformation of potato (*Solanum tuberosum*). In: Potrykus I, Spangenberg G editors, Gene Transfer to Plants. Springer-Verlag, Berlin; 1995. pp. 24–29.

28. Stiller I, Dancs G, Hesse H, Hoefgen R, Bánfalvi Z. Improving the nutritive value of tubers: Elevation of cysteine and glutathione contents in the potato cultivar ‘White Lady’ by marker-free transformation. J Biotechnol. 2007;128: 335–343.

29. Shure M, Wessler S, Fedoroff N. Molecular identification and isolation of the *Waxy* locus of maize. Cell. 1983;35: 225–233.

30. Stiekema WJ, Heidekamp F, Dirkse WG, van Beckum J, de Haan P, Bosch CT, et al. Molecular cloning and analysis of four potato tuber mRNAs. Plant Mol Biol. 1988;11: 255–269.

31. Cernák I, Taller J, Wolf I, Fehér E, Babinszky G, Alföldi Z, et al. Analysis of the applicability of molecular markers linked to the PVY extreme resistance gene *Ry*_*sto*_, and the identification of new markers. Acta Biol Hung. 2008;59: 195–203.

32. Decsi K, Cernák I, Bánfalvi Z, Korom E, Wolf I, Vaszily Z, et al. Marker assisted selection of the *Solanum stoloniferum* based PVY resistance in the breeding material of Keszthely. ScienceMED. 2012;3: 215–219.

33. Song Y-S, Schwarzfischer A. Development of STS markers for selection of extreme resistance (*Ry*_*sto*_) to PVY and maternal pedigree analysis of extremely resistant cultivars. Am J Potato Res. 2008;85: 159–170.

34. Sambrook J, Fritsch EF, Maniatis T. Molecular Cloning: A Laboratory Manual, 2^nd^ edn. Cold Spring Harbor, New York, Cold Spring Harbor Laboratory; 1989.

35. Coil D, Jospin G, Darling AE. A5-miseq: an updated pipeline to assemble microbial genomes from Illumina MiSeq data. Bioinformatics. 2015;31: 587–589.

36. Potato Genome Sequencing Consortium. Genome sequence and analysis of the tuber crop potato. Nature. 2011;475: 189–195.

37. Li H. Aligning sequence reads, clone sequences and assembly contigs with BWA-MEM. 2013;arXiv:1303.3997.

38. Bevan M. Binary *Agrobacterium* vectors for plant transformation. Nucleic Acids Res. 1984;12: 8711–8721.

39. Hanahan D. Studies on transformation of *Escherichia coli* with plasmids. J Mol Biol. 1983;166: 557–580.

40. Milbourne D, Meyer RC, Collins AJ, Ramsay LD, Gebhardt C, Waugh R. Isolation, characterization and mapping of simple sequence repeat loci in potato. Mol Gen Genet. 1998;259: 233–245.

41. The Tomato Genome Consortium. The tomato genome sequence provides insights into fleshy fruit evolution. Nature. 2012;485: 635–641.

42. Grech-Baran M, Witek K, Szajko K, Witek AI, Morgiewicz K, Wasilewicz-Flis I, et al. Extreme resistance to *Potato Virus Y* in potato carrying the *Ry*_*sto*_ gene is mediated by a TIR-NLR immune receptor. Plant Biotechnol J. 2019; https://doi.org/10.1111/pbi.13230

43. Ortega F, Lopez-Vizcon C. Application of molecular marker-assisted selection (MAS) for disease resistance in a practical potato breeding programme. Potato Res. 2012; 55:1–13

44. Fulladolsa AC, Navarro FM, Kota R, Severson K, Palta JP, Charkowski AO. Application of marker assisted selection for Potato Virus Y resistance in the University of Wisconsin Potato Breeding Program. Am J Potato Res. 2015;92: 444–450.

45. Nie X, Lalany F, Dickison V, Wilson D, Singh M, De Koeyer D, et al. Detection of molecular markers linked to *Ry* genes in potato germplasm for marker-assisted selection for extreme resistance to PVY in AAFC’s potato breeding program. Can J Plant Sci. 2016;96: 737–742.

46. Bakker E, Borm T, Prins P, van der Vossen E, Uenk G, Arens M, et al. A genome-wide genetic map of NB-LRR disease resistance loci in potato. Theor Appl Genet. 2011;123: 493–508.

47. van Eck HJ, Vos PG, Valkonen JPT, Uitdewilligen JGAML, Lensing H, de Vetten N, et al. Graphical genotyping as a method to map *Ny*_*(o,n)sto*_ and *Gpa5* using a reference panel of tetraploid potato cultivars. Theor Appl Genet. 2017;130: 515–528.

48. Poczai P, Cernák I, Gorji AM, Nagy S, Taller J, Polgár Z. Development of intron targeting (IT) markers for potato and cross-species amplification in *Solanum nigrum* (Solanaceae). Am J Bot. 2010;97: e142–145.

49. Jupe F, Witek K, Verweij W, Sliwka J, Pritchard L, Etherington GJ, et al. Resistance gene enrichment sequencing (RenSeq) enables reannotation of the *NB-LRR* gene family from sequenced plant genomes and rapid mapping of resistance loci in segregating populations. Plant J. 2013;76: 530–544.

50. McDowell JM, Dhandaydham M, Long TA, Aarts MGM, Goff S, Holub EB, et al. Intragenic recombination and diversifying selection contribute to the evolution of downy mildew resistance at the *RPP8* locus of arabidopsis. Plant Cell. 1998;10: 1861–1874.

51. Kehr J. Phloem sap proteins: their identities and potential roles in the interaction between plants and phloem-feeding insects. J Exp Bot. 2006;57: 767–774.

52. Whitham S, Dinesh-Kumar SP, Choi D, Hehl R, Corr C, Baker B. The product of the tobacco mosaic virus resistance gene *N*: Similarity to toll and the interleukin-1 receptor. Cell. 1994;78: 1101–1115.

53. Faize M, Faize L, Burgos L. Using quantitative real-time PCR to detect chimeras in transgenic tobacco and apricot and to monitor their dissociation. BMC Biotechnol. 2010;10: 53.

